# System drift and speciation

**DOI:** 10.1101/231209

**Authors:** Josh S Schiffman, Peter L Ralph

## Abstract

Even if a species’ phenotype does not change over evolutionary time, the underlying mechanism may change, as distinct molecular pathways can realize identical phenotypes. Here we use linear system theory to explore the consequences of this idea, describing how a gene network underlying a conserved phenotype evolves, as the genetic drift of small changes to these molecular pathways cause a population to explore the set of mechanisms with identical phenotypes. To do this, we model an organism’s internal state as a linear system of differential equations for which the environment provides input and the phenotype is the output, in which context there exists an exact characterization of the set of all mechanisms that give the same input–output relationship. This characterization implies that selectively neutral directions in genotype space should be common and that the evolutionary exploration of these distinct but equivalent mechanisms can lead to the reproductive incompatibility of independently evolving populations. This evolutionary exploration, or *system drift*, is expected to proceed at a rate proportional to the amount of intrapopulation genetic variation divided by the effective population size (*N*_*e*_). At biologically reasonable parameter values this could lead to substantial interpopulation incompatibility, and thus speciation, on a time scale of *N*_*e*_ generations. This model also naturally predicts Haldane’s rule, thus providing a concrete explanation of why heterogametic hybrids tend to be disrupted more often than homogametes during the early stages of speciation.

## Introduction

It is an overarching goal of many biological subdisciplines to attain a general understanding of the function and evolution of the complex molecular machinery that translates an organism’s genome into the characteristics on which natural selection acts, the phenotype. For example, there is a growing body of data on the evolutionary histories and molecular characterizations of particular gene regulatory networks [Jaeger, 2011, Davidson and Erwin, 2006, Israel et al., 2016], as well as thoughtful verbal and conceptual models [True and Haag, 2001, Weiss and Fullerton, 2000, Edelman and Gally, 2001, Pavlicev and Wagner, 2012]. Mathematical models of both particular regulatory networks and the evolution of such systems in general can provide guidance where intuition fails, and thus have the potential to discover general principles in the organization of biological systems as well as provide concrete numerical predictions [Servedio et al., 2014]. There is a substantial amount of work studying the evolution of gene regulatory networks, in frameworks both abstract [Wagner, 1994, 1996, Siegal and Bergman, 2002, Bergman and Siegal, 2003, Draghi and Whitlock, 2015] and empirically inspired [Mjolsness et al., 1991, Jaeger et al., 2004, Kozlov et al., 2015, Crombach et al., 2016, Wotton et al., 2015, Chertkova et al., 2017].

At all levels of biological organization, the problems that biological systems have evolved to solve often do not have single solutions – systems can be structurally different yet remain functionally equivalent [Edelman and Gally, 2001]. Examples can be found across nearly all levels of biological organization from the level of the genetic code itself all the way up to the convergent evolution of adaptive traits. In many cases, these functionally equivalent structures can be explored through small, local changes to the structure that leave the function unchanged. For instance, there are “neutral networks” of nucleic acid sequences that produce the same RNA secondary structure [Grüner et al., 1996] amino acid sequences that fold similarly [Babajide et al., 1997], or proteins with equivalent thermodynamic stability [Hart et al., 2014]. Further examples are found in the vast space of functionally equivalent potential regulatory sequences [Hare et al., 2008], in the logic of transcriptional [Tsong et al., 2006, Matsui et al., 2015, Dalal et al., 2016, Dalal and Johnson, 2017, Jiménez et al., 2017] and neural circuits [Trojanowski et al., 2014], and in developmental systems [von Dassow et al., 2000, True and Haag, 2001].

This capacity for isofunctional yet distinct mechanisms, sometimes called *degeneracy*, is a consequence of a many-to-one mapping between a system’s structure and function, a concept that has been explored in many fields beyond biology. For instance, in many contexts mathematical models are fundamentally *nonidentifiable* and/or *indistinguishable* – meaning that there can be uncertainty about an inferred model’s parameters or even its claims about causal structure, despite access to complete and perfect data [e.g., Bellman and Åström, 1970, Grewal and Glover, 1976, Walter et al., 1984]. Models with different parameter schemes, or even different mechanics can make equally accurate predictions, but still not actually reflect the internal dynamics of the system being modeled. In control theory, where electrical circuits and mechanical systems are often the focus, it is understood that there can be an infinite number of “realizations”, or ways to reverse engineer the dynamics of a “black box”, even if all possible input and output experiments are performed [Kalman, 1963, Anderson et al., 1966, Zadeh and Deoser, 1976]. The inherent nonidentifiability of chemical reaction networks is sometimes referred to as “the fundamental dogma of chemical kinetics” [Craciun and Pantea, 2008]. In computer science, this has been framed as the relationship among processes that *simulate* one another [Van der Schaft, 2004]. Finally, the field of *inverse problems* studies those cases in which, despite the existence of a theoretical one-to-one mapping between a model and behavior, tiny amounts of noise make inference problems nonidentifiable in practice [Petrov and Sizikov, 2005].

It has been argued that the ability to modify structure without affecting function is necessary for natural selection [Edelman and Gally, 2001], as it may function as a mechanism for biological robustness and evolvability [reviewed in de Visser et al., 2003], or manifest as *canalization* [Whitacre, 2010]. It may even contribute to the formation of new species [Gavrilets, 2014]. Redundancy of the genetic code, for instance, can make sequences more fault-tolerant to mutations [Sonneborn, 1965], and robustness to modification of genetic networks can allow adaptation without passing through a fitness valley [Wagner, 2008].

In this paper we use results on mathematical nonidentifiability from linear systems theory to study how gene regulatory networks can be modified while retaining the same function, and the possible implications for speciation. If system architectures are not functionally unique, can this open up neutral evolutionary paths, and do species explore these paths through the process termed *developmental system drift* [by True and Haag, 2001]? Is this fast enough to contribute meaningfully to speciation? To do this, we describe results on linear dynamical systems which give an analytical description of the set of all linear gene network architectures that yield identical phenotypes, and use quantitative genetics theory to estimate the speed at which system drift can lead to reproductive incompatibility and hence speciation. In this model, a population diffuses along the neutral ridges of a high-dimensional space of possible system parameters, in a similar vein as *holey landscape* models [Gavrilets, 1997, Yamaguchi and Iwasa, 2013].

The field of population genetics has also explored the consequences of the fact that there is often more than one way to do the same thing, and observed that speciation might be the result of changes that are themselves neutral. Indeed, Bateson [1909] first proposed that what today we call a Bateson-Dobzhansky-Muller incompatibility would likely arise through neutral changes. The potential for speciation has been analyzed in models of traits under stabilizing selection determined additively by alleles at many loci [Wright, 1935, Barton, 1986, 1989, 2001], in related fitness landscape models [Fraïsse et al., 2016], and for pairs of traits that must match but whose value is unconstrained [Sved, 1981]. It has also been shown that population structure can allow long-term stable coexistence of incompatible genotypes encoding identical phenotypes [Phillips, 1996]. However, previous simulations of system drift in regulatory sequences [Tulchinsky et al., 2014] and a regulatory cascade [Porter and Johnson, 2002] found rapid speciation under directional selection but only equivocal support for speciation under models of purely neutral drift. The rate at which hybrid incompatibility accumulates due to genetic drift creating segregation variance between isolated populations is fairly well understood [Slatkin and Lande, 1994, Rosas et al., 2010, Chevin et al., 2014], but model assumptions can strongly affect predictions, including whether variation is due to rare or common alleles [Slatkin and Lande, 1994], and the shape of the fitness landscape [Fraïsse et al., 2016]. Our main aim is to provide a concrete framework that can make natural predictions of these model parameters across a general class of models. Furthermore, tools from system theory allow analytical predictions to be made for large populations with complex phenotypes that would be inaccessible to population simulations.

## Results

We use a model of gene regulatory networks that describes the temporal dynamics of a collection of *n* coregulating molecules – such as transcription factors – as well as external or environmental inputs. We write *κ*(*t*) for the vector of *n* molecular concentrations at time *t*. The vector of *m* “inputs” determined exogenously to the system is denoted *u*(*t*), and the vector of *ℓ* “outputs” is denoted *ϕ*(*t*). The output is merely a linear function of the internal state: *ϕ*_*i*_(*t*) = Σ_*j*_ *C*_*ij*_*κ*_*j*_(*t*) for some matrix *C*. Since *ϕ* is what natural selection acts on, we refer to it as the *phenotype* (meaning the “visible” aspects of the organism), and in contrast refer to *κ* as the *kryptotype*, as it is “hidden” from direct selection. Although *ϕ* may depend on all entries of *κ*, it is usually of lower dimension than *κ*, and we tend to think of it as the subset of molecules relevant for survival. The dynamics are determined by the matrix of regulatory coefficients, *A*, a time-varying vector of inputs *u*(*t*), and a matrix *B* that encodes the effect of each entry of *u* on the elements of the kryptotype. The rate at which the *i*^th^ concentration changes is a weighted sum of the concentrations as well as the input:

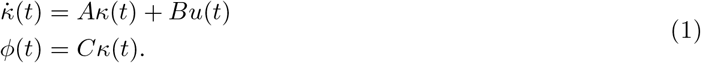

Furthermore, we always assume that *κ*(0) = 0, so that the kryptotype measures deviations from initial concentrations. Here *A* can be any *n* × *n* matrix, *B* any *n* × *m*, and *C* any *ℓ* × *n* dimensional matrix, with usually *ℓ* and *m* less than *n*. We think of the system as the triple (*A, B, C*), which translates (time-varying) *m*-dimensional input *u*(*t*) into the *ℓ*-dimensional output *ϕ*(*t*). Under quite general assumptions on the input (e.g., |*u*(*t*)| is integrable) we can write the phenotype as

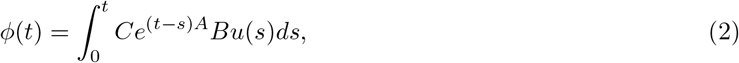

which is a convolution of the input *u*(*t*) with the system’s *impulse response*, which we denote as *h*(*t*) := *Ce*^*At*^*B*.

In terms of gene regulatory networks, *A*_*ij*_ determines how the *j*^th^ transcription factor regulates the *i*^th^ transcription factor. If *A*_*ij*_ *>* 0, then *κ*_*j*_ upregulates *κ*_*i*_, while if *A*_*ij*_ *<* 0, then *κ*_*j*_ downregulates *κ*_*i*_. The *i*^th^ row of *A* is therefore determined by genetic features such as the strength of *j*-binding sites in the promoter of gene *i*, factors affecting chromatin accessibility near gene *i*, or basal transcription machinery activity. The form of *B* determines how the environment influences transcription factor expression levels, and *C* might determine the rate of production of downstream enzymes.

Wagner [1994] and others have used a similar discrete-time model (that might be written *ϕ*_*t*+1_ = *f* (*Aϕ*_*t*_), where *f* is a sigmoid). Our choice of continuous time does not affect the points we make here, but our restriction to *linear* systems is a stronger assumption (see the Discussion).

To demonstrate the model, we construct a simple gene network in Example 1 below.

### Example 1

(An oscillator). *For illustration, we consider an extremely simplified model of oscillating gene transcription, as for instance is found in cell cycle control or the circadian rhythm. There are two genes, whose transcript concentrations are given by κ*_1_(*t*) *and κ*_2_(*t*), *and gene-2 upregulates gene-1, while gene-1 downregulates gene-2 with equal strength. Only the dynamics of gene-1 are consequential to the oscillator (perhaps the amount of gene-1 activates another downstream gene network). Lastly, both genes are equally upregulated by an exogenous signal. The dynamics of the system are described by*

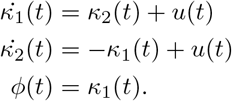

*In matrix form the system regulatory coefficients are given as*, 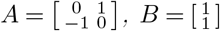, *and C* = [1 0]. *If the input is an impulse at time zero (a delta function), then the phenotype is equal to the impulse response:*

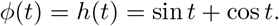

*The system and its dynamics are referred to in Figure 1. We return to the evolution of such a system below*.

**Figure 1:**
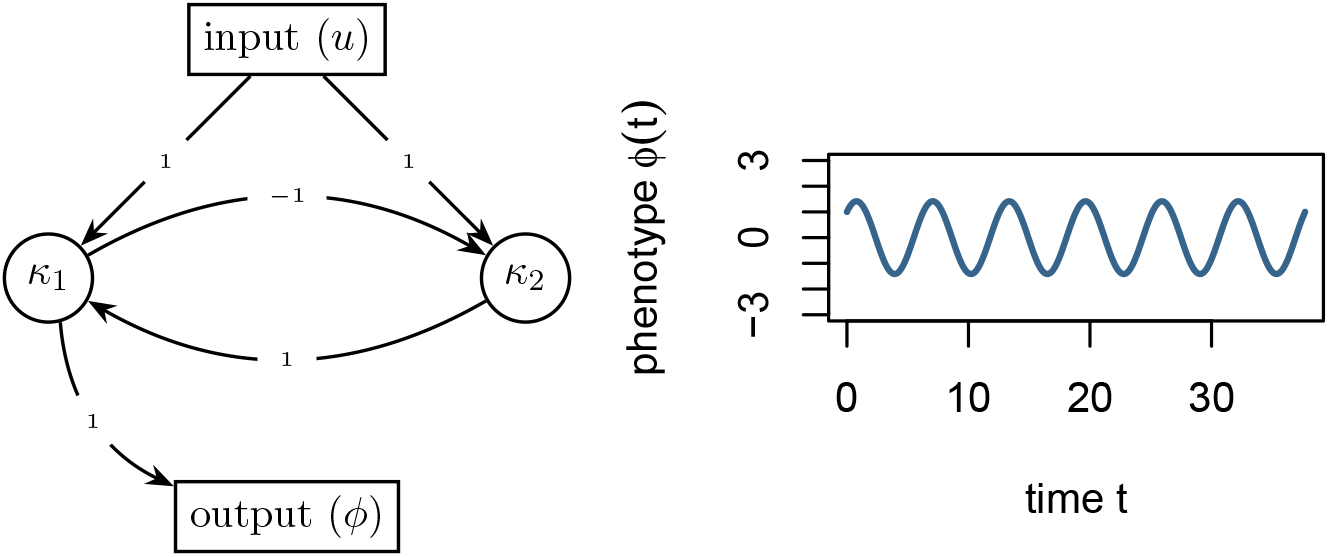
(Left) Diagram of the gene network in Example 1, and (right) plot of the phenotype *ϕ*(*t*) against time *t*.

### Equivalent gene networks

As reviewed above, some systems with identical phenotypes are known to differ, sometimes substantially, at the molecular level; systems with identical phenotypes do not necessarily have identical kryptotypes. How many different mechanisms perform the same function?

Two systems are equivalent if they produce the same phenotype given the same input, i.e., have the same input–output relationship. We say that the systems defined by (*A, B, C*) and 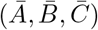 are **phenotypically equivalent** if their impulse response functions are the same: 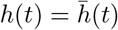 for all *t* ≥ 0. This implies that for any acceptable input *u*(*t*), if (*κ*_*u*_(*t*), *ϕ*_*u*_(*t*)) and 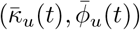 are the solutions to equation (1) of these two systems, respectively, then

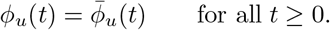

In other words, phenotypically equivalent systems respond identically for *any* input.

One way to find other systems phenotypically equivalent to a given one is by change of coordinates: if *V* is an invertible matrix, then the systems (*A, B, C*) and (*V AV* ^−1^, *V B, CV* ^−1^) are phenotypically equivalent because their impulse response functions are equal:

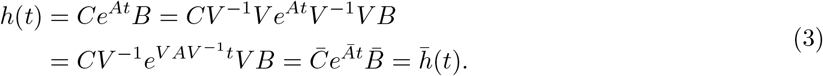

These “changes of coordinates” are not simply different ways of looking at the same system – if each dimension of the kryptotype corresponds to the concentration of a particular transcription factor, changing *A* corresponds to changing the strengths of regulatory interactions. We will even see below that interactions may change sign. However, not all phenotypically equivalent systems are of this form: systems can have identical impulse responses without being coordinate changes of each other. In fact, systems with identical impulse responses can involve interactions between different numbers of molecules, and thus have kryptotypes in different dimensions altogether.

This implies that most systems have at least *n*^2^ degrees of freedom, where recall *n* is the number of components of the kryptotype vector. This is because for an arbitrary *n* × *n* matrix *Z*, taking *V* to be the identity matrix plus a small perturbation in the direction of *Z* above implies that moving *A* in the direction of *ZA* − *AZ* while also moving *B* in the direction of *ZB* and *C* in the direction of −*CZ* will leave the phenotype unchanged to second order in the size of the perturbation. If the columns of *B* and the rows of *C* are not all eigenvectors of *A*, then any such *Z* will result in a different system.

It turns out that in general, there are more degrees of freedom, except if the system is *minimal* – meaning, informally, that it uses the smallest possible number of components to achieve the desired dynamics. Results in system theory show that any time-invariant system can be realized in a particular minimal dimension (the dimension of the kryptotype, *n*_min_), and that any two phenotypically equivalent systems of dimension *n*_min_ are related by a change of coordinates. Since gene networks can grow or shrink following gene duplications and deletions, these additional degrees of freedom can apply, in principle, to any system.

Even if the system is not minimal, results from systems theory explicitly describe the set of all phenotypically equivalent systems. We refer to *𝒩* (*A*_0_, *B*_0_, *C*_0_) as the set of all systems phenotypically equivalent to the system defined by (*A*_0_, *B*_0_, *C*_0_):

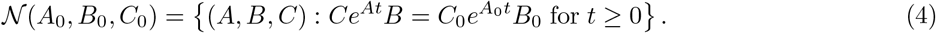

These systems need not have the same kryptotypic dimension *n*, but must have the same input and output dimensions (*ℓ* and *m*, respectively).

The Kalman decomposition, which we now describe informally, elegantly characterizes this set [Kalman, 1963, Kalman et al., 1969, Anderson et al., 1966]. To motivate this, first note that the input *u*(*t*) only directly pushes the system in certain directions (those lying in the span of the columns of *B*). As a result, different combinations of input can move the system in any direction that lies in what is known as the *reachable subspace*. Analogously, we can only observe motion of the system in certain directions (those lying in the span of the rows of *C*), and so can only infer motion in what is known as the *observable subspace*. The Kalman decomposition then classifies each direction in kryptotype space as either reachable or unreachable, and as either observable or unobservable. Only the components that are both reachable and observable determine the system’s phenotype – that is, components that both respond to an input and produce an observable output.

Concretely, the **Kalman decomposition** of a system (*A, B, C*) gives a change of basis *P* such that the transformed system (*PAP* ^−1^, *PB, CP* ^−1^) can be written in block matrix form:

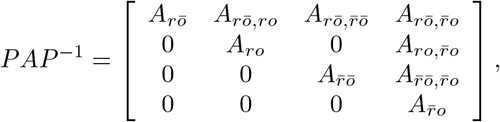

and

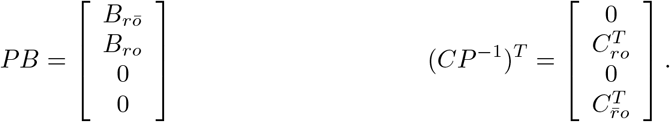

The *n*-dimensional system has been divided into subspaces of dimensions 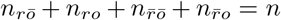, and so, for instance, *A*_*rō*_ is the *n*_*rō*_ *n*_*rō*_ square matrix in the top left corner of *PAP* ^−1^. The impulse response of the system is given by

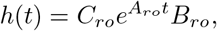

and therefore, the system is phenotypically equivalent to the *minimal* system (*A*_*ro*_, *B*_*ro*_, *C*_*ro*_).

This decomposition is unique up to a change of basis that preserves the block structure. In particular, the minimal subsystem obtained by the Kalman decomposition is unique up to a change of coordinates. This implies that there is no equivalent system with a smaller number of kryptotypic dimensions than the dimension of the minimal system. It is remarkable that the gene regulatory network architecture to achieve a given input–output map is never unique – both the change of basis used to obtain the decomposition and, once in this form, all submatrices other than *A*_*ro*_, *B*_*ro*_, and *C*_*ro*_ can be changed without affecting the phenotype, and so represent degrees of freedom.

Although the other submatrices do not affect the phenotype through the impulse response function *h*(*t*), they may affect individual fitness in other ways (which we don’t consider here). For instance, the “unreachable” subspaces cannot be affected by any of the possible variations in input produced by varying *u*(*t*). However, if the nature of the input changed (for instance, by perturbing a previously constant environmental variable), differences in the unreachable components might lead to differences in phenotypes and thus represent cryptic genetic variation. Similarly, “unobservable” subspaces might represent aspects of phenotype that are not constrained by selection, but might become important if conditions change. Additionally, all subspaces may affect how the system deals with noise.

#### Note on implementation

The *reachable subspace* is defined to be the closure of span(*B*) under applying *A* (or equivalently, the span of *B, AB, A*^2^*B*, … *A*^*n*−1^*B*), and the *unobservable subspace* is the largest *A*-invariant subspace contained in the null space of *C*. The four subspaces, 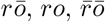, and 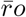 are defined from these by intersections and orthogonal complements – *ro* refers to the both *reachable and observable* subspace, while 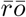 refers to the *unreachable and unobservable* subspace, and similarly for 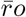 and *rō*.

For the remainder of the paper, we interpret 𝒩 as the neutral set in the fitness landscape, along which a large population will drift under environmental and selective stasis. This drift need not be purely neutral – for instance, second-order selection on robustness will push the species towards “flatter” areas of genotype space [Rice, 1998, Hermisson et al., 2003]. Even if the phenotype is constrained and remains constant through evolutionary time, the molecular mechanism underpinning it is not constrained and likely will not be conserved.

Finally, note that if *B* and *C* are held constant – i.e., if the relationships between environment, kryptotype, and phenotype do not change – there are *still* usually degrees of freedom. Example 2 gives the set of minimal systems equivalent to the oscillator of Example 1, that all share common *B* and *C* matrices. The oscillator can also be equivalently realized by a three-gene (or larger) network, and will have even more evolutionary degrees of freedom available, as in Figure 3.

##### Example 2

(All equivalent rewirings of the oscillator). *The oscillator of Example 1 is minimal, and so any equivalent system is a change of coordinates by an invertible matrix V*. *If we further require B and C to be invariant then we need V B* = *B and CV* = *C. Therefore the following one-parameter family* (*A*(*τ*), *B, C*) *describes the set of all two-gene systems phenotypically equivalent to the oscillator:*

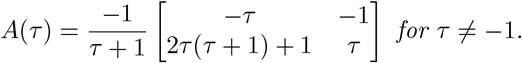

*The resulting set of systems are depicted in Figure 2*.

**Figure 2:**
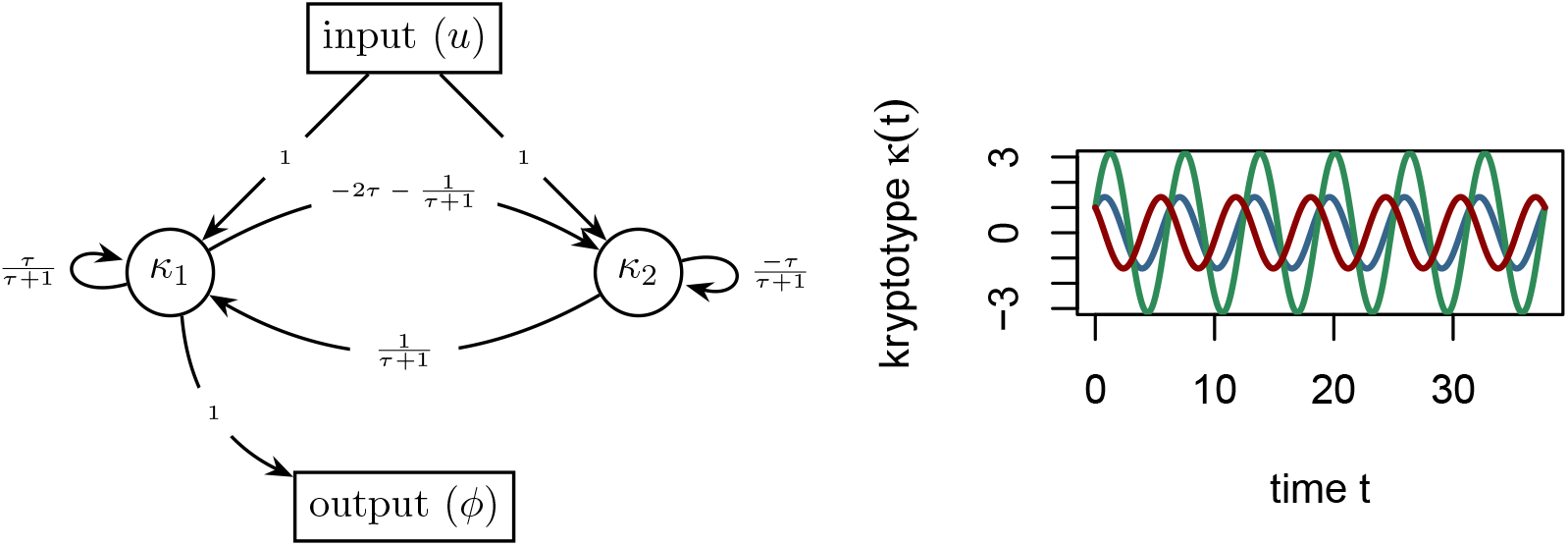
(Left) *A*(*τ*), the set of all phenotype-equivalent cell cycle control networks. (Right) Gene-1 dynamics (blue) for both systems *A*(0) and *A*(−2) are identical, however, *A*(0) gene-2 dynamics (red) differ from *A*(−2) (green).

**Figure 3:**
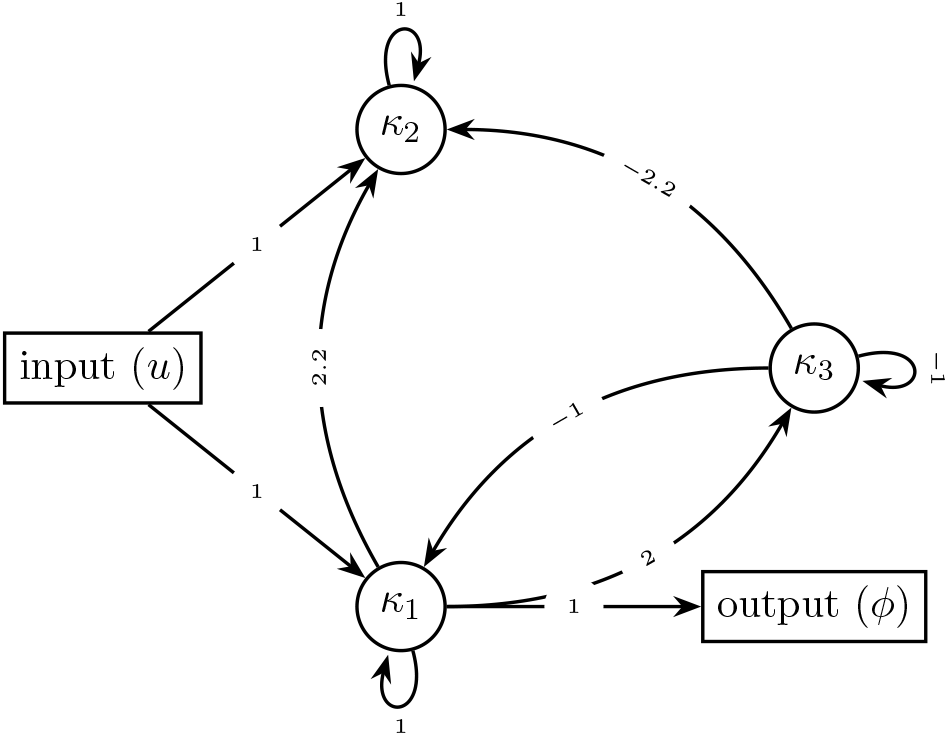
A possible non-minimal three-gene oscillator, phenotypically equivalent to *A*(*τ*), the systems in Examples 1 and 2.

### Sexual reproduction and recombination

Parents with phenotypically equivalent yet differently wired gene networks may produce offspring with dramatically different phenotypes. If the phenotypes are significantly divergent then the offspring may be inviable or otherwise dysfunctional, despite both parents being well adapted. If this is consistent for the entire population, we would consider them to be separate species, in accord with the biological species concept [Mayr, 2000].

First, we must specify how sexual reproduction acts on these systems. Suppose that each of a diploid organisms’ two genomes encodes a set of system coefficients with the same kryptotype dimension. We assume that a diploid which has inherited systems (*A′, B′, C′*) and (*A ″, B″, C″*) from its two parents has phenotype determined by the system that averages these two, ((*A′* + *A″*)*/*2, (*B′* + *B″*)*/*2, (*C′* + *C″*)*/*2). In general, how parental systems are recombined and inherited depends on the genetic architecture of the segregating variation. For the purposes of examples in this paper (Example 3 and Figure 5), we take the simplest assumption that each system coefficient is encoded by a nonrecombining locus, and that these loci are unlinked. For instance, if one genome’s coefficients are written with Greek and the other with Latin, a possible oscillator gamete might be produced as follows: 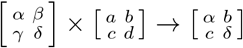.

Each genome an organism inherits is generated by meiosis, in which both of its diploid parents recombine their two genomes, and so an *F*_1_ offspring carries one system copy from each parent, and an *F*_2_ is an offspring of two independently formed *F*_1_s. If the parents are from distinct populations, these are simply first– and second–generation hybrids, respectively.

Exactly how the coefficients (i.e., entries of *A, B* and *C*) of a haploid system inherited by an offspring from a diploid parent are determined by the parent’s two systems depends on the genetic basis of any variation in the coefficients. Thanks to the randomness of meiotic segregation, the result is random to the extent that each parent is heterozygous for alleles that affect the coefficients. Since the *i*^th^ row of *A* summarizes how each gene regulates gene *i*, and hence is determined by the promoter region of gene *i*, the elements of a row of *A* tend to be inherited together, which will create covariance between entries of the same row. It is, however, a quite general observation that the variation seen among recombinant systems is proportional to the difference between the two parental systems.

Offspring formed from two phenotypically identical systems do not necessarily exhibit the same phenotype as both of its parents – in other words 𝒩, the set of all systems phenotypically equivalent to a given one, is not, in general, closed under averaging or recombination. If sexual recombination among systems drawn from 𝒩 yields systems with divergent phenotypes, populations containing significant diversity in 𝒩 can carry genetic load, and isolated populations may fail to produce hybrids with viable phenotypes.

### Hybrid incompatibility

Two parents with the optimal phenotype can produce offspring whose phenotype is suboptimal if the parents have different underlying systems. Hybrid phenotypic break down, as a function of genetic distance between phenotypically equivalent parental oscillators (described in Example 2) is illustrated in Example 3. How quickly do hybrid phenotypes break down as genetic distance between parents increases? We will quantify how far a system’s phenotype is from optimal using a weighted difference between impulse response functions. Suppose that *ρ*(*t*) is a nonnegative weighting function, *h*_0_(*t*) is the *optimal* impulse response function and define the “distance to optimum” of another impulse response function to be

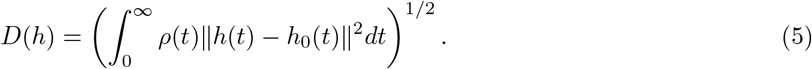

In practice, we take *ρ*(*t*) = exp(−*t/*4*π*), so that fitness is determined by the dynamics of the system over a few multiples of 2*π*, but not longer. Consider reproduction between a parent with system (*A, B, C*) and another displaced by distance *ϵ* in the direction (*X, Y, Z*), i.e., having system (*A* + *ϵX, B* + *ϵY, C* + *ϵZ*). We assume both are “perfectly adapted” systems, i.e., having impulse response function *h*_0_(*t*), and their offspring has impulse response function *h*_*ϵ*_ (*t*). A Taylor expansion of *D*(*h*_*ϵ*_) in *E* is explicitly worked out in Appendix A, and shows that the phenotype of an *F*_1_ hybrid between these two is at distance proportional to *ϵ*^2^ from optimal, while *F*_2_ hybrids are at distance proportional to *ϵ*. This is because an *F*_1_ hybrid has one copy of each parental system, and therefore lies directly between the parental systems (see Figure 4) – the parents both lie in *𝒩*, which is the valley defined by *D*, and so their midpoint only differs from optimal due to curvature of *𝒩*. In contrast, an *F*_2_ hybrid may be homozygous for one parental type in some coefficients and homozygous for the other parental type in others; this means that each coefficient of an *F*_2_ may be equal to either one of the parents, or intermediate between the two; this means that possible *F*_2_ systems may be as far from the optimal set, *𝒩*, as the distance between the parents. The precise rate at which the phenotype of a hybrid diverges depends on the geometry of the optimal set *𝒩* relative to segregating genetic variation.

**Figure 4:**
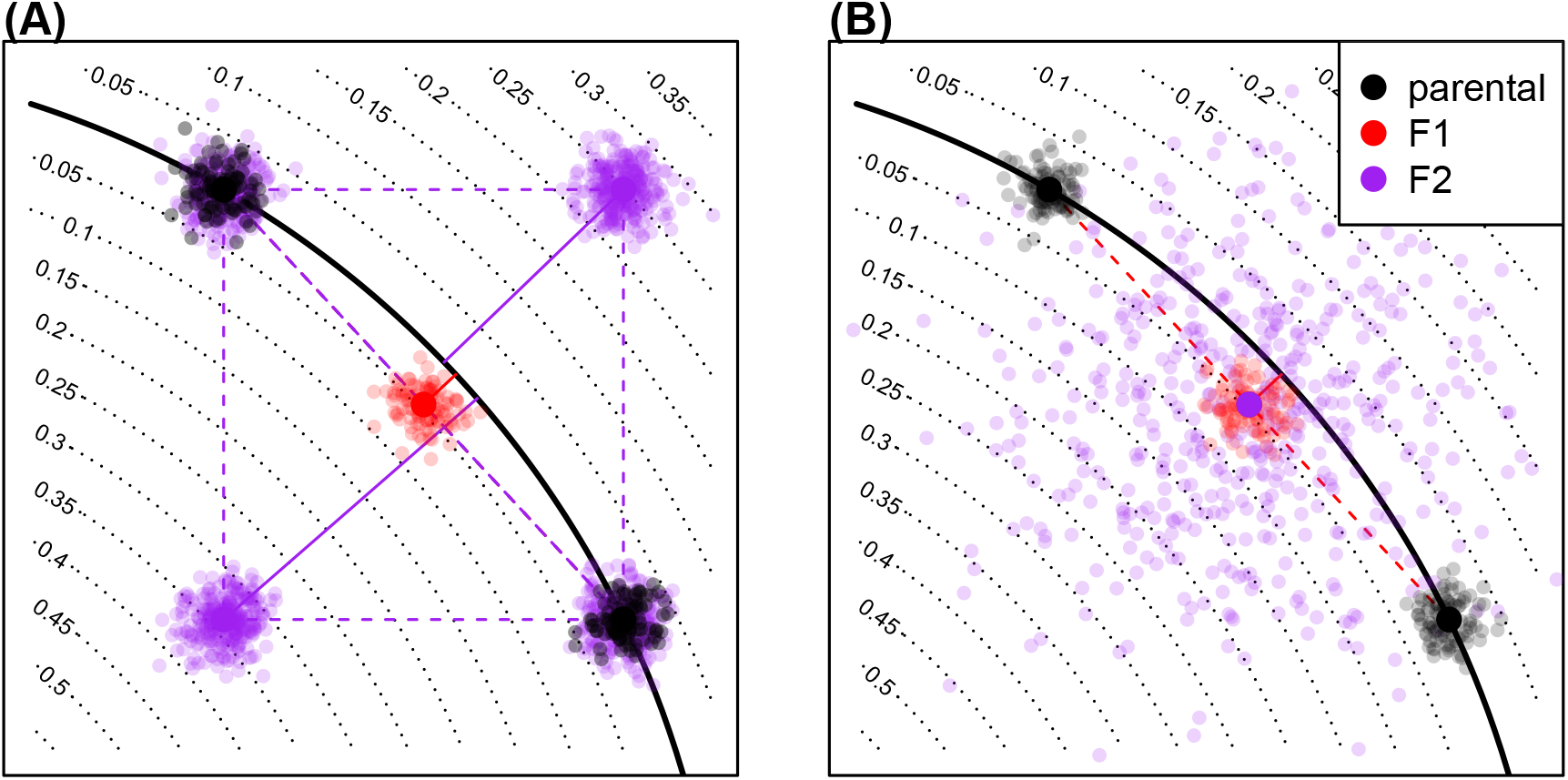
A conceptual figure of the fitness consequences of hybridization: axes represent system coefficients (i.e., entries of *A*); the line of optimal system coefficients is down in black; solid lines give phenotypic distances to the optimum. A pair of parental populations are shown in black, along the optimum; a hypothetical population of *F*_1_s are shown in red, and the distribution of *F*_2_s is shown in purple. The two figures differ in the genetic basis, and hence, the distribution of *F*_2_ phenotypes: **(A)** *F*_2_s compose all four mixed homozygotes if variation at both traits has a simple, one-locus genetic basis in both populations; and **(B)** *F*_2_ show a much wider distribution of phenotypes if the genetic basis of variation in each population is polygenic.

#### Example 3

(Hybrid incompatibility: misregulation due to system drift). *Offspring of two equivalent systems from Example 2 can easily fail to oscillate. For instance, the F*_1_ *offspring between homozygous parents at τ* = 0 *and τ* = −2 *has phenotype* 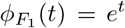, *rather than ϕ*(*t*) = sin *t* + cos *t. However, the coefficients of these two parental systems differ substantially, probably more than would be observed between diverging populations. In Figure 5 we compare the phenotypes for F*_1_ *and F*_2_ *hybrids between more similar parents, and see increasingly divergent phenotypes as the difference between the parental systems increases. (In this example, the coefficients of A*(*ϵ*) *differ from those of A*(0) *by an average factor of* 1 + *ϵ/*2; *such small differences could plausibly be caused by changes to promoter sequences*.*) This divergence is quantified in Figure 6, which shows that mean distance to optimum phenotype of the F*_1_ *and F*_2_ *hybrid offspring between A*(0) *and A*(*ϵ*) *increases with ϵ*^2^ *and ϵ, respectively*.

**Figure 5:**
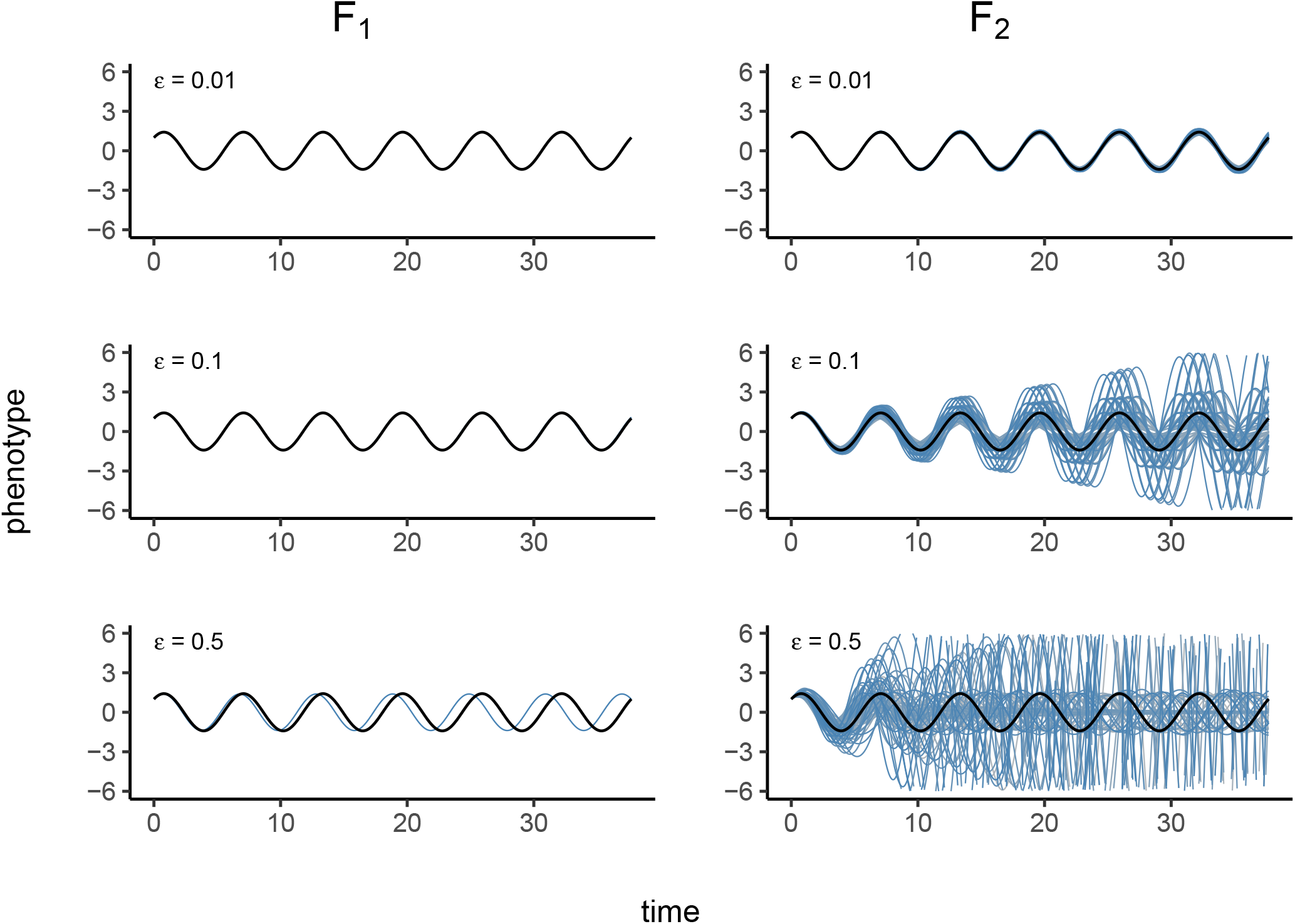
**(left)** Phenotypes of *F*_1_ hybrids between a homozygous *A*(0) parent and, top-to-bottom, homozygous *A*(1*/*100), *A*(1*/*10), and *A*(1*/*2) parents, where *A*(*ϵ*) is defined in Figure 2; parental coefficients differ by around 0.5%, 5%, and 25% respectively. Parental phenotypes (sin *t* + cos *t*) are shown in solid black, and hybrid phenotypes in blue. **(right)** Phenotypes of all 3^4^ = 81 possible *F*_2_ hybrids between the same set of parents, with parental phenotype again in black. *F*_2_ hybrids, many of which show complete breakdown, are shown in shades of blue and grey.

**Figure 6:**
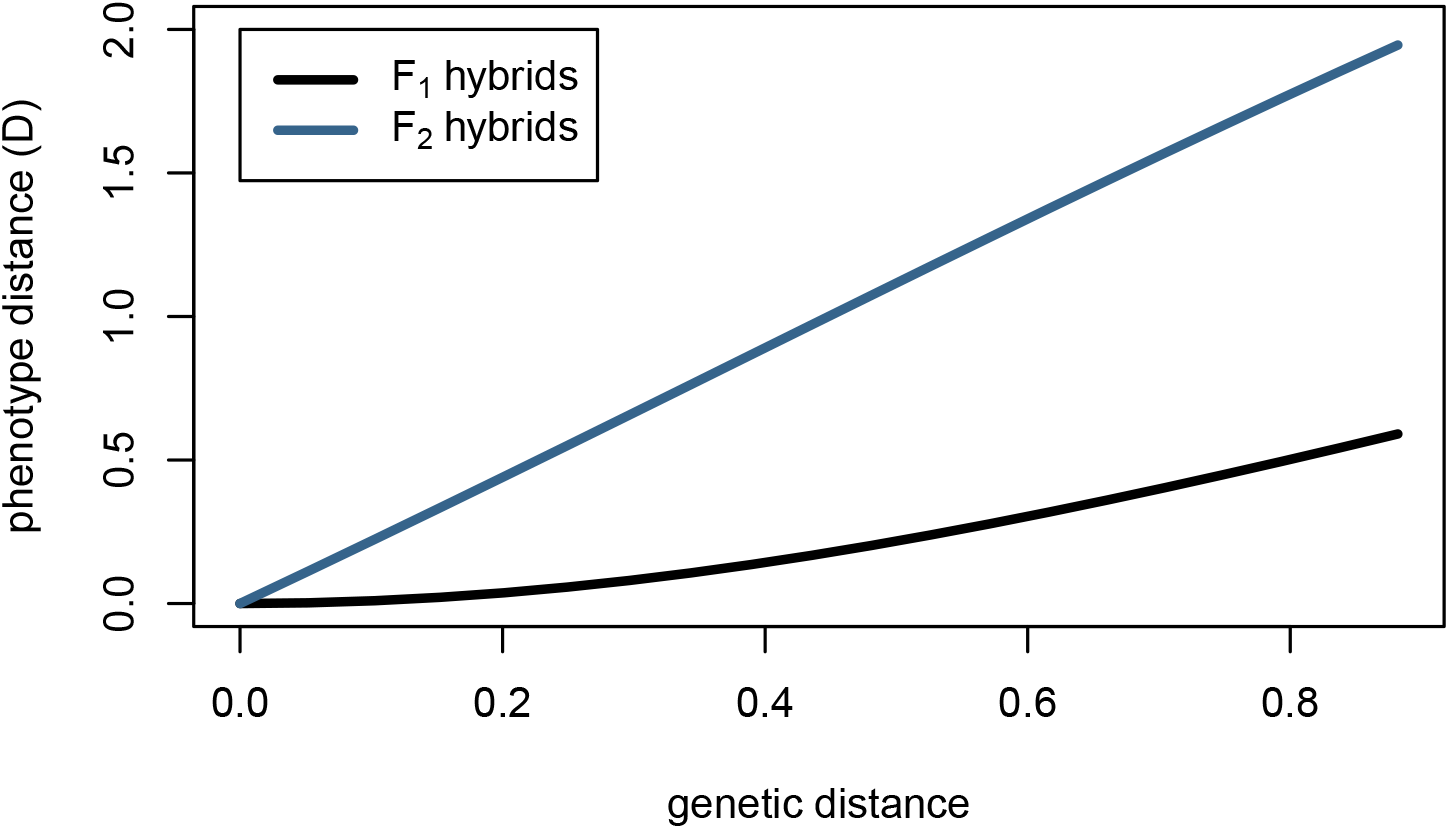
Mean hybrid phenotypic distance from optimum computed with equation (5), using *ρ*(*t*) = exp(−*t/*4*π*) for *F*_1_ (black) and *F*_2_ (blue) hybrids between *A*(0) and *A*(*ϵ*) parent oscillators. Genetic distance is computed as (Σ_*ij*_ (*A*_*ij*_(0) − *A*_*ij*_(*ϵ*))^2^) ^½^

### Haldane’s rule

This model naturally predicts Haldane’s rule, the observation that if only one hybrid sex is sterile or inviable it is likely the heterogametic sex (e.g., the male in XY sex determination systems) [Haldane, 1922, Orr, 1997]. For example, consider an XY species with a two-gene network where the first gene resides on an autosome and the second gene on the X chromosome. A male whose pair of haplotypes is 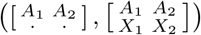 has phenotype determined by 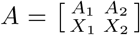, if dosage compensation upregulates heterogametes by a factor of two relative to homogametes (as with *Drosophila*), while a female homozygous for the haplotype 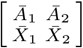, has phenotype determined by 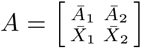. An *F*_1_ male offspring of these two will have its phenotype determined by 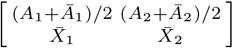. If both genes resided on the autosomes, this system would only be possible in an *F*_2_ cross. More generally, the contribution of sex chromosomes to the system coefficients of *F*_1_ males are effectively equivalent to those of purely autosomal *F*_2_ hybrids, which are less fit on average than *F*_1_s (see Figure 6). Note, however, that presence of genetic variation on sex chromosomes does not alone guarantee Haldane’s rule: under this model, Haldane’s rule will only occur when there is sufficient genetic distance separating parentals and when the nature of genetic variation produces *F*_2_ hybrid breakdown.

### The speed of speciation

We have shown that system drift can lead to speciation in principle, but is it rapid enough to be an important factor in practice? In other words, after what period of time would we expect the fitness of hybrids between two allopatric populations to be substantially lower than the parentals? Selection – on pleiotropic traits or on robustness – may actively push even a strongly constrained system along neutral directions, but even the calculations under purely neutral drift are informative. The population mean of an unconstrained quantitative trait with additive genetic variance *V*_*G*_ in a population with effective size *N*_*e*_ will move in *t* generations a random amount whose variance is *tV*_*G*_*/N*_*e*_ [Lande, 1976]. The mean difference between two such populations has twice the variance. Although this mean difference is along neutral directions, we would in many cases expect the range of variation among *F*_2_s in *all* directions to be of the same order as the differences between the populations, as depicted in Figure 4. This suggests that, naively, two such populations that have been separated for *t* generations will produce *F*_2_ offspring that differ from optimal by an amount proportional to 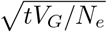. Since we assume they are at a local fitness optimum, without much loss of generality we can assume that fitness is locally quadratic, and so *F*_2_ fitness decays linearly in time: proportionally to *tV*_*G*_*/N*_*e*_ – fastest in small, diverse populations, This predicts that we need only wait some multiple of *N*_*e*_ generations until substantial incompatibility has been accumulated.

It is useful to think in more detail about the assumptions in the rough argument above. The key aspect is how population differences in neutral directions (along the fitness ridge) translate to segregation variance in *F*_2_s in selectively constrained directions. To move the system (the *A* matrix) a given distance generally involves moving many individual interaction coefficients (the entries *A*_*ij*_). The movements must be coordinated, for the population to stay near the fitness ridge. However, mixing elements between systems that have made independent sets of coordinated changes to remain on the fitness ridge is unlikely to produce a set of coordinated changes; and the resulting system could move away from the ridge in almost any direction.

### Genetic variation in empirical regulatory systems

What is known about the key quantity above, the amount of heritable variation in real regulatory networks? The coefficient *A*_*ij*_ from the system (1) measures how much the rate of net production of *i* changes per change in concentration of *j*. It is generally thought that regulatory sequence change contributes much more to inter- and intraspecific variation than does coding sequence change affecting molecular structure [Schmidt et al., 2010]. In the context of transcription factor networks this may be affected not only by the binding strength of molecule *j* to the promoter region of gene *i* but also the effects of other transcription factors (e.g., cooperativity) and local chromatin accessibility [Stefflova et al., 2013]. For this reason, the mutational target size for variation in *A*_*ij*_ may be much larger than the dozens of base pairs typically implicated in the handful of binding sites for transcription factor *j* of a typical promoter region, and single variants may affect many entries of 𝒩 simultaneously.

Variation in binding site occupancy may overestimate variation in *A*, since it does not capture buffering effects (if for instance only one site of many needs to be occupied for transcription to begin), and variation in expression level measures changes in steady-state concentration (our *κ*_*i*_) rather than the *rate* of change. Nonetheless, these measures likely give us an idea of the scale of variability. It has been shown that between human individuals, there is differential occupancy in 7.5% of binding sites of a transcription factor (p65) [Kasowski et al., 2010]. It has also been inferred that cis-regulatory variation accounts for around 2–6% of expression variation in human blood-derived primary cells [Verlaan et al., 2009], and that human population variation explained about 3% of expression variation [Lappalainen et al., 2013]. Allele-specific expression is indicative of standing genetic *cis*-regulatory variation; allele-specific expression in 7.2–8.5% of transcripts of a flycatcher species has been observed [Wang et al., 2017], as well as allele-specific expression in 23.4% of genes studied in a baboon species [Tung et al., 2015]. Taken together, this suggests that variation in the entries of *A* may be on the order of at least a few percent between individuals of a population – doubtless varying substantially between species and between genes.

## Discussion

In this paper, we use tools from linear system theory and quantitative genetics to study the evolution of a mechanistic model of the genotype-phenotype map, in which the phenotype is subject to stabilizing selection. In so doing, we provide an explicit model of phenogenetic drift [Weiss and Fullerton, 2000] and developmental system drift [True and Haag, 2001]. In this context, the Kalman decomposition [Kalman, 1963] gives an analytical description of all phenotypically equivalent gene networks. This description shows that the space of functionally equivalent network architectures increases with the square of a network’s size, and that this space increases further if networks grow larger than absolutely necessary – that is use more interacting components than the most efficient potential architectures. In this framework, even minimal gene network architectures – efficient architectures that contain only the requisite number of interacting parts, are not structurally unique with respect to function. Functionally equivalent architectures are often related by continuous parameter changes, suggesting that equivalent networks might be mutationally connected, and that there exist axes of genetic variation unconstrained by natural selection. The independent movement of separated populations along these axes by genetic drift can lead to a significant reduction in hybrid viability, and thus precipitate speciation, at a speed dependent on the effective population size and the amount of genetic variation. In this model, at biologically reasonable parameter values, system drift is a significant – and possibly rapid – driver of speciation. This may be surprising because hybrid inviability appears as a consequence of recombining different, yet functionally equivalent, mechanisms, and since species are often defined by their unique adaptations or morphologies.

Consistent with empirical observation of hybrid breakdown we see that the fitnesses of *F*_2_ hybrids drop at a much faster rate than those of *F*_1_s. Another natural consequence of the model is Haldane’s rule, that if only one *F*_1_ hybrid sex is inviable or sterile it is likely to be the heterogametic sex. This occurs because if the genes underlying a regulatory network are distributed among both autosomes and the sex chromosome, then heterogametic *F*_1_s show variation (and fitnesses) similar to that seen in *F*_2_ hybrids. This observation appears to be similar to the extreme hybrid phenotypes produced by transgressive segregation [Rieseberg et al., 1999], which can manifest in *F*_1_s when only one (dominant) parental allele is expressed at heterozygous loci; this was observed in hybrid gene expression patterns, and increased as a function of parental genetic distance [Stelkens and Seehausen, 2009].

Is there evidence that this is actually occurring? System drift and network rewiring has been inferred across the tree of life [Wotton et al., 2015, Crombach et al., 2016, Dalal and Johnson, 2017, Johnson, 2017, Ali et al., 2019], and there is often significant regulatory variation segregating within populations. Transcription in hybrids between closely related species with conserved transcriptional patterns can also be divergent [Haerty and Singh, 2006, Maheshwari and Barbash, 2012, Coolon et al., 2014, Michalak and Noor, 2004, Mack and Nachman, 2016], and hybrid incompatibilities have been attributed to cryptic molecular divergence underlying conserved body plans [Gavin-Smyth and Matute, 2013]. Furthermore, in cryptic species complexes (e.g., sun skinks [Barley et al., 2013]), genetically distinct species may be nearly morphologically indistinguishable.

### The origin of species not by means of natural selection?

As classically formulated, the Dobzhansky-Muller model of hybrid incompatibility is agnostic to the relative importance of neutral versus selected genetic substitutions [Coyne and Orr, 1998], and plausible mechanisms have been proposed whereby Dobzhansky– Muller incompatibilities could originate under neutral genetic drift [Lynch and Force, 2000] or stabilizing selection [Fierst and Hansen, 2009]. The same holds for the “pathway model” [Lindtke and Buerkle, 2015], which is closer to the situation here. However, previous authors have argued that neutral processes are likely too slow to be a significant driver of speciation [Nei et al., 1983, Seehausen et al., 2014]. This has led some to conclude that hybrid incompatibility is typically a byproduct of positive selection [Orr et al., 2004, Schluter, 2009] or a consequence of genetic conflict [Presgraves, 2010, Crespi and Nosil, 2013], two processes that typically act much more rapidly than genetic drift. Supporting this view, experimental approaches designed to study the effect of drift on speciation through induced population bottlenecks found no reproductive isolation in the majority of tested pairs [Fry, White et al., 2020]. However, our calculations suggest that even under strictly neutral processes, hybrid fitness breaks down as a function of genetic distance rapidly enough to play a substantial role in species formation across the tree of life. This is consistent with broad patterns such as the relationship between molecular divergence and genetic isolation seen by Roux et al. [2016], and the clocklike speciation rates observed by Hedges et al. [2015]. More recently, Sánchez-Ramírez et al. [2021] observed widespread transcriptional misregulation in *Drosophila* hybrids, which might be due to the action of system drift.

Although the phrase “system drift” as we use it implies neutral changes, it does not mean that these changes are induced purely by neutral genetic drift. Selection could dramatically accelerate speciation by the same process – e.g., by genetic draft [Gillespie, 2000] or pleiotropic selection. For instance, Mani and Clarke [1990] showed that populations developed incompatibilities as they independently evolved towards the same quantitative trait optimum, as different alleles fix in different populations. Fluctuating selection has similarly been shown to accelerate speciation in some models [Bell, 2010, Barton, 2001]. Under our model, populations evolving towards different optimal phenotypes will always produce less fit hybrids, as the hybrid phenotypes will be displaced from both parental optima, and so maladapted to both parental environments. This suggests that fluctuating selection – if not synchronized between populations – should diminish hybrid fitnesses. In the case where fluctuations are synchronized, due to the size of the neutral system space, populations may arrive at different system organizations when returning to previous phenotypes at a rate faster than by drift alone.

These explanations are not mutually exclusive. All of these forces – adaptive shifts, conflict and network drift – are plausible drivers of speciation, and may even interact. Many of our observations carry over to models of directional selection – for instance, rapid drift along the set of equivalent systems could be driven by adaptation in a different, pleiotropically coupled system. Or, reinforcement due to local adaptation might provide a selective pressure that speeds up system drift. Furthermore, while the fitness consequences of incompatibility in any one given network may be small, the cumulative impact of system drift across the many different networks an organism relies on may be substantial. It remains to be seen how the relative strengths of these forces compare.

### The dimensionality of trait space

We have focused on examples of single traits (where the phenotype is one-dimensional), but phenotypes under selection are often high-dimensional, and variation in different traits often share a genetic basis. However, we still expect many degrees of freedom as long as there are components of the system not directly and individually constrained by selection (i.e., a kryptotype). Even in networks where the phenotype and kryptotype are of the same dimension, system theory shows us that there will always be available degrees of freedom as specific system realizations are only unique up to a change of coordinates. Some phenotypes, however, require kryptotypic dimensions to be larger than that of the phenotype. For instance, many systems have minimal realizations (*e*.*g*., the oscillator in Example 2) where the dimension of the kryptotype is larger than that of the phenotype, implying that for these phenotypic dynamics to be realized, the kryptotype dimension *has* to be larger than the dimension of the phenotype. Even if components of the system’s internal state are directly subject to selection and the mode of action of the environment on the internal state is constrained (so, the input and output matrices *B* and *C* are fixed) then one could still perturb *A* as described above by *ZA* − *AZ* if *ZB* and *CZ* are both zero, implying a number of degrees of freedom that still grows with *n*^2^ for fixed *ℓ* and *m*. Generically, the number of degrees of freedom is *n*(*n* − *ℓ* − *m*), so that in a system of *n* components, if even one component is not directly constrained, this leads to *n* degrees of freedom. Whatever the true “dimensionality” of phenotype space of a typical organism, there are undoubtedly aspects of its underlying molecular machinery that are not directly constrained, suggesting large numbers of degrees of freedom. Note that pleiotropy does not directly affect this argument at all – indeed, many phenotypically equivalent changes will lead to denser *A* matrices and hence more pleiotropy. However, more pleiotropic genes may be more strongly constrained, making it more difficult for systems to make the required compensatory changes for system drift.

Phenotypically equivalent system evolution is probably not only driven by neutral genetic drift. For one thing, movement along the set of equivalent networks is not expected to be completely neutral, since second-order selection pushes populations towards “flatter” regions of the fitness landscape in which a population centered on the optimal set has lower genetic load [as described in different contexts by Rice, 1998, Nimwegen et al., 1999]. If this bias towards more robust networks is strong enough, it may even prevent drift, but it is unclear how strong this effect would be in practice. Our results, on the other hand, do not rely on the flatness of the fitness surface around the phenotypically equivalent set, but rather on the curvature of the equivalent set itself. So long as the phenotypically equivalent set is not closed under sexual recombination, opportunities for incompatibility remain. However the speed at which system drift can generate incompatibilities might diminish if selection for robustness is strong enough to constrain a population to a small section of system space, although the strength of such effects in practice are not known. Likewise, as the speed of system drift relies on segregating genetic variation, any constraints on such variation, possibly due to epistasis, genetic architecture [Hermisson et al., 2003], adaptive inertia [Baatz and Wagner, 1997, Álvarez-Castro et al., 2009] or weak gene flow could plausibly slow it down. More work on specific systems, likely coupled with simulations, will be necessary to identify the biologically relevant parameter regimes.

### Geometric models

The model we work with describes a continuum of possible systems, parameterized by continuous coefficient matrices. The arguments we make (e.g., Figure 4) are often geometric in nature, and so it would be interesting to ask whether Fisher’s geometric model might provide more quantitative, generalizable predictions, as in for instance Fraïsse et al. [2016], Simon et al. [2017]. Martin [2014] argued that such an approximation would be often fruitful. In the other direction, it is also natural to ask about the distribution of effect sizes of segregating mutations in a population evolving under this model with a realistic *de novo* effect size distribution, as well as what the distribution of fitness effects would be (both within and between populations). These are questions that might be best answered by simulation, and could provide much additional information – for instance, are alleles typically dominant in their effects on either system coefficients or fitness?

### Nonlinearity and model assumptions

Of course, real regulatory networks are not linear dynamical systems. Most notably, physiological limits put upper bounds on expression levels, implying saturating response curves. It remains to be seen how well our results carry over into real systems, but the fact that most nonlinear systems can be locally approximated by a linear one suggests our qualitative results may hold more generally. Furthermore, nonidentifiability (which implies the existence of neutral directions) is often found in practice in moderately complex models of biological systems [e.g., Gutenkunst et al., 2007, Piazza et al., 2008, Jiménez et al., 2017].

Finally, despite our model’s precise separation of phenotype and kryptotype, this relationship in nature may be far more complicated as aspects of the kryptotype may be less “hidden” than we currently assume, and the neutral network changes we describe here may only be nearly neutral. For instance, attributes excluded from the phenotype as modeled here ignore the potential energy costs associated with excessively large (non-minimal) kryptotypes, as well as the relationship between a specific network architecture and robustness to mutational, transcriptional, or environmental noise. More precise modeling will require better mechanistic understanding not only of biological systems, but also the nature of selective pressures and genetic variation in these systems.

## Acknowledgements

We would like to thank Sergey Nuzhdin, Stevan Arnold, Michael Turelli, Patrick Phillips, Erik Lundgren and Hossein Asgharian for valuable discussion. We would also like to thank Nick Barton, Sarah Signor, Todd Parsons, and Joachim Hermisson for very helpful comments on the manuscript. Work on this project was supported by funds from the Sloan Foundation and the NSF (under DBI-1262645) to PR.

## A Local expansion of the fitness surface

The fitness of a system depends on the difference between the system’s impulse response and the optimal impulse response, measured as a weighted sum of the distance between the impulse response from optimal. With *ρ*(*t*) ≥ 0 a weighting function on [0, ∞), and 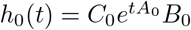 a representative of the optimal set, the distance is equation (5):

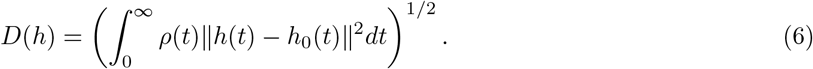

To see how this is affected by small changes to the system, first note that if || *·* ||_*ρ*_ is the *L*^2^(*ρ*) norm,

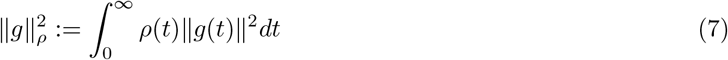

then the distance to optimum of a perturbed system, *h*(*t*) = *h*_0_(*t*) + *ϵg*(*t*), is

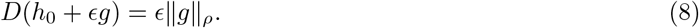

Now consider the perturbed systems (*A*(*ϵ*), *B*(*ϵ*), *C*(*ϵ*)) = (*A*(0), *B*(0), *C*(0)) + *ϵ* (*U, V, W*), and let *h*_*ϵ*_ (*t*) *C*(*ϵ*)*e*^*A*(*ϵ*)*t*^*B*(*ϵ*). If two populations with systems (*A*(0), *B*(0), *C*(0)) and (*A*(*ϵ*), *B*(*ϵ*), *C*(*ϵ*)) interbreed, then the mean resulting system is (*A*(*ϵ/*2), *B*(*ϵ/*2), *C*(*ϵ/*2)), but with greater variance in the *F*_2_ than the *F*_1_ offspring (see Figure 4). Indeed, if the difference between (say) *A*(0)_*ij*_ and *A*(*ϵ*)_*ij*_ is due to *n* fixed differences with absolute additive effect size 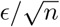 each, then the standard deviation of *A*_*ij*_ among the *F*_1_ offspring is still equal to that within each parent population, as each locus is heterozygous. However, if loci are unlinked then an *F*_2_ offspring is homozygous with probability 1/2, and takes either homozygote with equal probability, so the variance of the contribution of each locus is 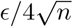, and so the standard deviation across *F*_2_ offspring is *ϵ/*4 plus a contribution for within-population genetic variance. (This many differences of this size would be expected if the difference was due to drift, for instance.) This implies that the impulse response, *h*(*t*), of an *F*_2_ will also differ from optimum by something of order *ϵ*. However, since the displacement between the populations lies along a ridge, the *F*_1_ offspring are closer: the Taylor expansions for *h* about both 0 and *ϵ* are

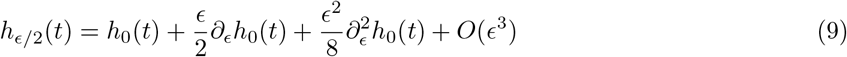

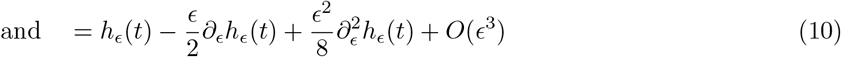

Since *h*_*ϵ*_ (*t*) = *h*_0_(*t*) by assumption, and 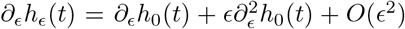 combining these expressions we get that

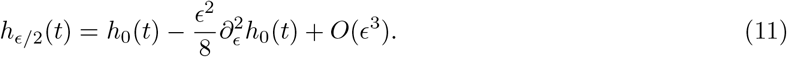

In other words, if the parental populations are both optimal but differ by *ϵ*, then the *F*_1_ offspring differ from optimal by something of order *𝒩*^2^, but the *F*_2_ by something of order *𝒩*.

Of course, when moving away from the ridge, the fitness landscape is flatter in some directions than others, and how much so depends on the details. For completeness, we describe here how to compute the sensitivity of distance to optimum with respect to each of *A, B*, and *C*. Changing notation slightly, let’s rewrite the squared distance to optimum of a system (*A, B, C*):

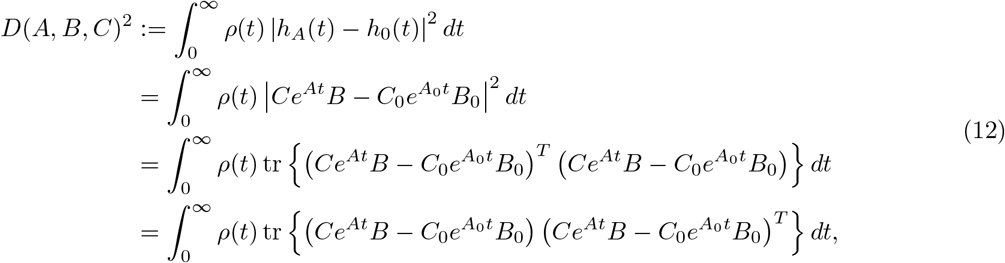

where tr *X* denotes the trace of a square matrix *X*. How does this change as we perturb about (*A*_0_, *B*_0_, *C*_0_)? First we differentiate with respect to *A*, keeping *B* = *B*_0_ and *C* = *C*_0_ fixed. Since

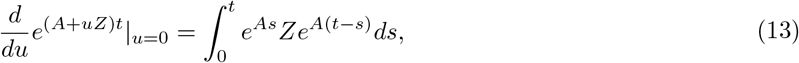

we have that

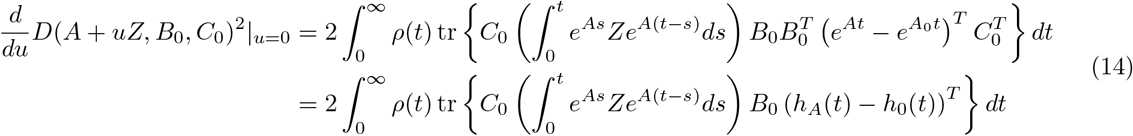

and, by differentiating this and supposing that *A* is on the optimal set, i.e., *h*_*A*_(*t*) = *h*_0_(*t*), (so without loss of generality, *A* = *A*_0_):

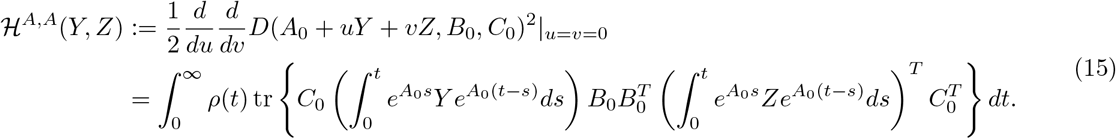

The function *ℋ* will define a quadratic form. To illustrate the use of this, suppose that *B* and *C* are fixed. By defining Δ_*ij*_ to be the matrix with a 1 in the (*i, j*)th slot and 0 elsewhere, the coefficients of the quadratic form are

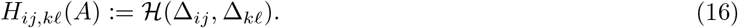

We could use this to get the quadratic approximation to *D* near the optimal set. To do so, it’d be nice to have a way to compute the inner integral above. Suppose that we diagonalize *A* = *U* Λ*U* ^−1^. Then

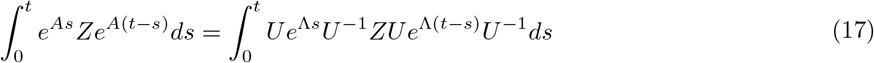

Now, notice that

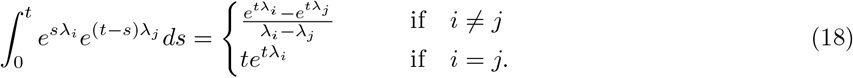

Therefore, defining

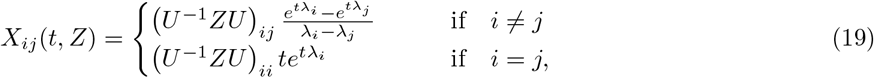

moving the *U* and *U* ^−1^ outside the integral and integrating we get that

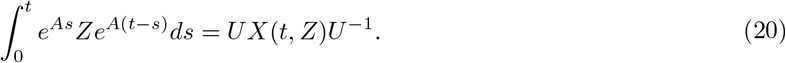

This implies that

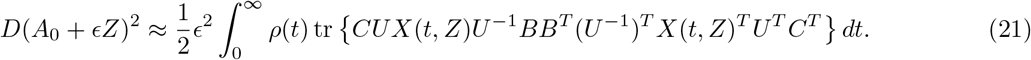

To compute the *n*^2^ *× n*^2^ matrix *H*, we see that if *Z* = Δ_*kℓ*_, then

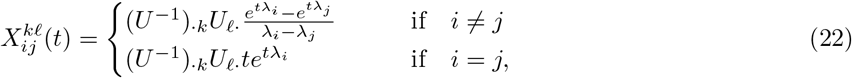

where *U*_*k·*_ is the *k*th row of *U*, and so

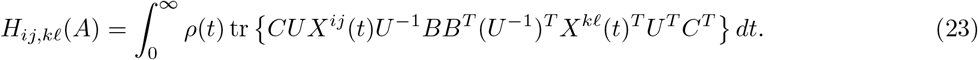

This implies that

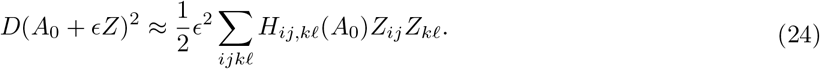

More generally, *B* and *C* may also change. To extend this we need the remaining second derivatives of *D*^2^. First, in *B*:

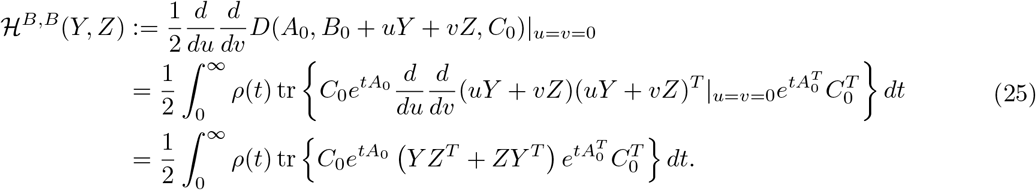

Next, in *C*:

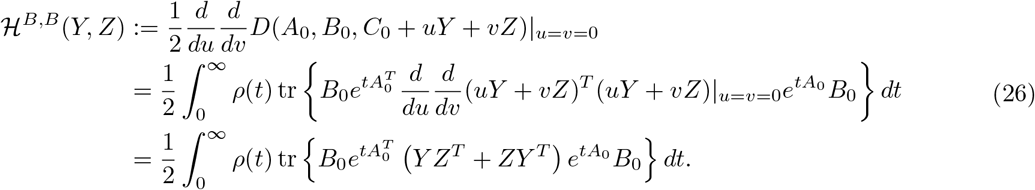

Now, the mixed derivatives in *B* and *C*:

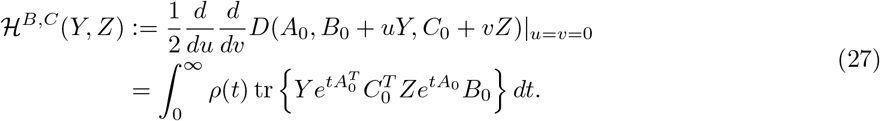

In *A* and *B*

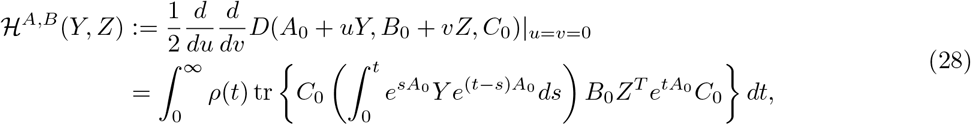

and finally in *A* and *C*:

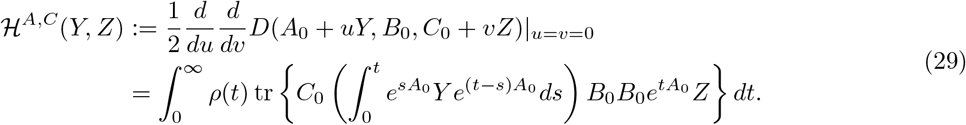

Together, numerical computation of these expressions, along with estimates of genetic covariance within a population, allow precise predictions of evolutionary dynamics of a particular system. The approximation should be good as long as the second-order Taylor approximation holds.

## Notes

### Competing Interest Statement

The authors have declared no competing interest.

### Summary of Updates

The section on quantitative genetics has been removed (and is destined for another paper).

